# Gold Nanocages as Scattering Contrast Agents for Optical Coherence Tomography

**DOI:** 10.1101/2025.03.14.643380

**Authors:** Yongping Chen, Jiefeng Xi, Du Lee, Jessica Ramella-Roman, Xingde Li

**Affiliations:** Department of Biomedical Engineering, Johns Hopkins University, Baltimore, Maryland 21205, USA; Biomedical Engineering Department, Florida International University, Miami, Florida 33174, USA

**Keywords:** Gold nanocage, Contrast agent, Optical coherence tomography (OCT), Scattering enhancement, Integrating sphere, Tumor imaging, Surface plasmon resonance, PEGylation, Nanoparticles

## Abstract

We report a scattering-dominant agent that significantly enhances optical coherence tomography (OCT) imaging contrast. The agent is based on gold nanocages, which exhibit a surface plasmon resonance (SPR) peak around 780 nm and a scattering cross-section that exceeds the absorption cross-section. The synthesis protocol for these nanocages is provided in detail. The optical properties of the gold nanocages were characterized using OCT imaging and further validated with integrating sphere measurements. OCT contrast enhancement was demonstrated through imaging of tissue phantoms containing embedded nanocages, as well as *ex vivo* imaging of mouse tissues and *in vivo* imaging of a mouse tumor following intravenous administration of PEGylated gold nanocages. To the best of our knowledge, this may be the first demonstration of scattering-dominant OCT contrast agents utilizing structured gold nanoparticles.

## INTRODUCTION

Optical coherence tomography (OCT) is a powerful technology capable of noninvasive and high-resolution imaging of biological tissues in real time.^1^ The contrast in OCT images is primarily determined by the intrinsic optical scattering and absorption properties of the biological tissues, and it is often governed by scattering. In many imaging modalities—such as X-ray computed tomography (CT), ultrasound (US), and magnetic resonance imaging (MRI)—the use of exogenous contrast agents is crucial to enhance image contrast.^2-5^ With no exception the use of exogenous contrast agents could potentially enhance OCT imaging capability. Various contrast agents, including core-shell microspheres,^6^ air-filled microbubbles,^7^ dyes,^8^ and structured gold nanoparticles,^9-11^ have been developed to enhance OCT image contrast. Among these, gold nanoparticles are particularly attractive due to their bioinertness, small size, large and customizable absorption and scattering cross-sections, and tunable surface plasmon resonance (SPR) peak wavelength. Ideally, scattering-dominant gold nanoparticles are preferred, as they can provide stronger backscattering contrast for OCT imaging. While gold nanospheres or nanoshells can, in principle, offer greater scattering than absorption, the size of these nanoparticles must be quite large (e.g., 150-200 nm in diameter) to achieve an SPR peak wavelength suitable for OCT, typically within the near-infrared range.

Gold nanocages are a recently developed class of structured nanoparticles with hollow interiors and ultrathin, porous walls. They are synthesized through a galvanic replacement reaction between silver (Ag) nanocubes (used as sacrificial templates) and HAuCl4 aqueous solution.^12–14^ By modulating the size, thickness, and porosity of the walls of the nanocages, their optical properties can be finely tuned. Compared to most other gold nanoparticles, gold nanocages generally exhibit stronger scattering and absorption in the near-infrared (NIR) region while maintaining a relatively small size (i.e., 100 nm or less), which is crucial for effective tissue delivery. These characteristics make gold nanocages particularly well-suited as contrast agents for OCT imaging.

Previously, our group demonstrated that gold nanocages with an SPR peak wavelength around 800 nm and a large absorption-dominant extinction cross-section could serve as an excellent agent for enhancing OCT spectroscopic contrast by preferentially attenuating certain wavelengths within the OCT source spectrum.^11, 15^ Considering that the OCT signal also depends on the sample’s backscattering and total scattering properties, OCT imaging contrast enhancement can also be achieved by using highly scattering agents. Unlike absorption, which only attenuates the OCT signal, an agent with a large scattering and backscattering cross-section could potentially enhance the backscattered signal from the sample. Thus, it has been highly desirable to develop gold nanocages with strong scattering dominance to enhance OCT imaging contrast.

In the present study, for the first time to our knowledge, gold nanocages with a strong scattering-dominant cross-section were successfully synthesized. The optical properties of these nanoparticles were characterized using a tissue phantom. In addition, the optical properties of the nanocages were also directly measured using a well-established integrating sphere method.^16^ Real-time OCT imaging was performed on *ex vivo* tissues and an *in vivo* mouse tumor model, demonstrating significant contrast enhancement by the scattering-dominant gold nanocages.

## RESULTS AND DISCUSSION

Gold nanocages were synthesized following the fundamental principle previously published,^17^ but with a significantly lower reaction temperature and slower titration rate of the reaction solution. Specifically, HAuCl4 aqueous solution was titrated into a solution of silver nanocubes (∼68 nm edge length, as shown in the inset of Figure 1A), which served as a sacrificial template at room temperature, with a titration rate of 0.25 mL/min. The silver templates were gradually etched away, resulting in gold nanocages with hollow interiors and porous walls. The optical properties, such as scattering, absorption, and SPR peak wavelength, of the gold nanocages can be precisely tuned by controlling the size, wall thickness, and porosity. Compared to previous gold nanocages reported for absorption-dominant 800 nm OCT imaging,^11, 15^ the gold nanocages used in this study, which exhibited a scattering-dominant cross-section, were less porous and had a slightly larger edge length. Figure 1A shows a transmission electron microscope (TEM) image of gold nanocages with a fairly uniform size distribution (∼75 ± 5.2 nm), and Figure 1B shows the UV-vis-NIR extinction spectrum of the gold nanocages around 780 nm.

**Figure 1.**
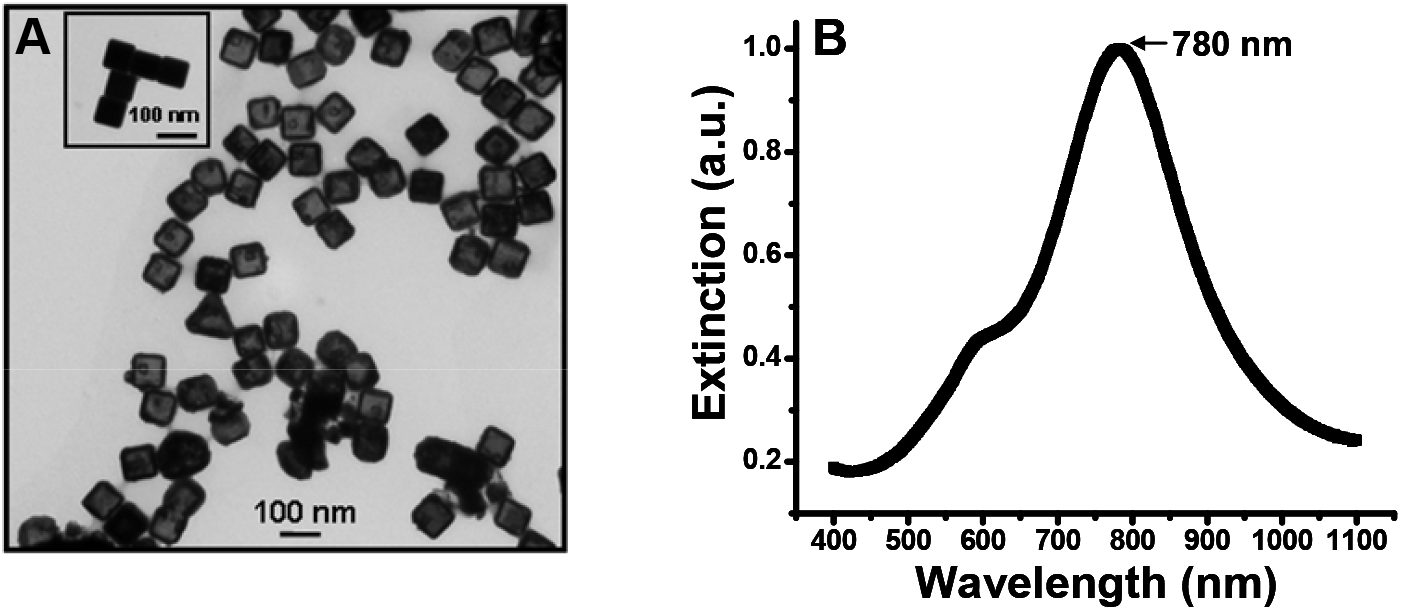
(A) TEM image of gold nanocages with an average edge length of 75 nm. The inset shows a TEM image of the Ag nanocubes with an average edge length of 68 nm. (B) UV-vis-NIR spectra of the gold nanocages with an SPR peak around 780 nm.

To quantitatively evaluate the optical properties of gold nanocages, we performed OCT imaging on tissue phantoms with and without gold nanocages. Two phantoms were made of 5% gelatin embedded with 50 mg/mL silica nanospheres as standard samples, whose optical properties can be predicted using Mie theory. Gold nanocages were added to one of the phantoms at a final concentration of 1 nM. Both phantoms were imaged by an 830 nm Spectral-domain OCT (SD-OCT) system under the same experimental conditions (e.g., the reference DC level, incident power on the sample, etc.), and the details on the SD-OCT system were reported previously.^18-20^ Figure 2A shows the OCT images of tissue phantoms without (on the left) and with (on the right) gold nanocages, respectively. The decay curves of the backscattered intensity along the imaging depth in both cases are shown and compared in Figure 2B. The results revealed an ∼33% increase in the peak of the backscattered signal from the phantom with gold nanocages compared to the one without nanocages.

**Figure 2.**
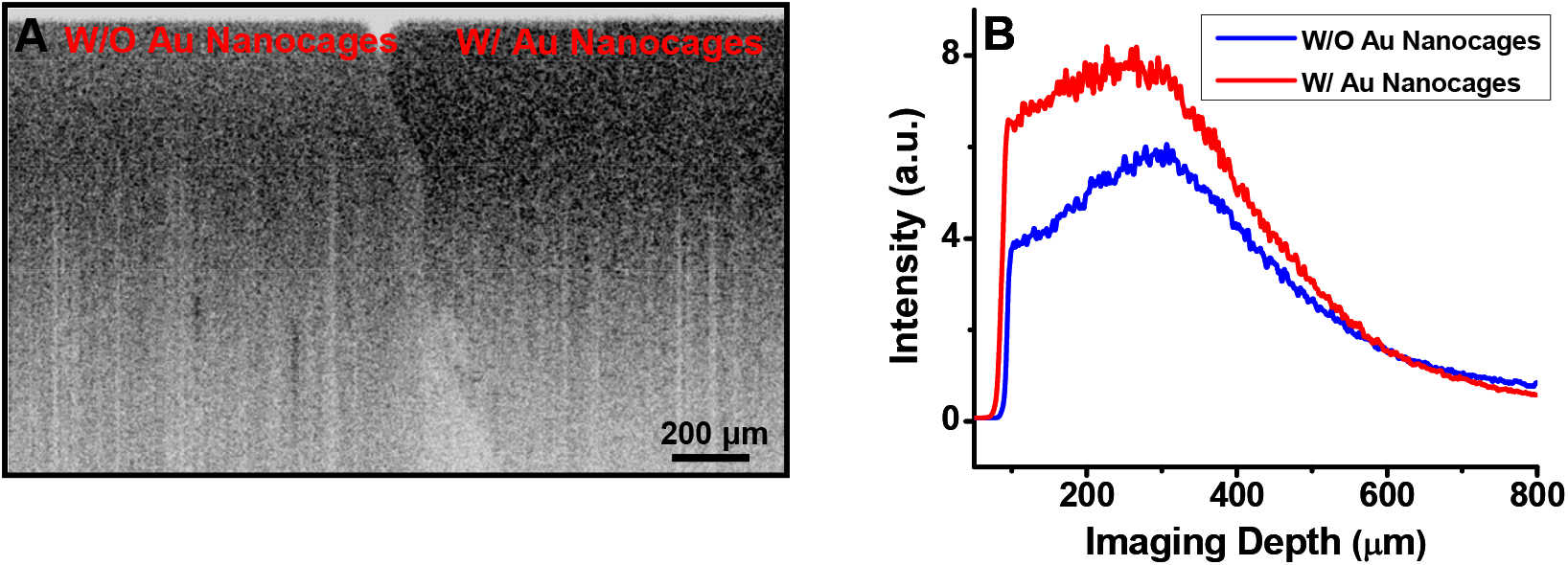
(A) OCT images of the phantoms without nanocages (left) and with nanocages (right). (B) Intensity plots of the OCT signals on a linear scale as a function of imaging depth. The strong backscattered signal from the phantom with gold nanocages is evident.

To quantify the optical extinction cross-sections of the gold nanocages, we first calculated the optical properties of silica nanospheres in the phantoms, including scattering and backscattering cross-sections, using Mie theory.^21^ Since both phantoms were imaged under the same conditions, we divided the depth-dependent OCT decay curve from the nanocage-embedded phantom by that from the phantom without nanocages to eliminate the influence of common factors (such as detector amplification and beam focusing effects) on the decay curve. The logarithm of the quotient was then fitted with a simple linear function, with two variables: the extinction coefficient of the nanocages and their relative backscattering coefficient with respect to silica nanospheres, from which the (absolute) backscattering coefficient of the nanocages was deduced. Assuming the angular scattering distribution of the gold nanocages does not vary much due to the cubic shape of the nanocages and their small size (relative to the incident wavelength), the (total) scattering coefficient of the gold nanocages was derived from the measured backscattering coefficient. Once the extinction coefficient *μ*_*t*_ and scattering coefficient *μ*_*s*_ were determined, the absorption coefficient of the nanocages *μ*_*a*_ was given by the difference between the two coefficients (i.e., *μ*_*t*_ -*μ*_*s*_). More details about the quantification process are provided in the Methods section.

Following the above analysis, we found the scattering and absorption coefficients of the nanocages to be 0.97 mm^-1^ and 0.74 mm^-1^, respectively, from which the ratio of the scattering cross-section C_sca_ to the absorption cross-section C_abs_ was found to be 1.31. To independently validate the optical properties obtained from OCT phantom imaging, integrating sphere experiments were performed to directly measure the optical properties of the nanocages. The ratio of C_sca_ to C_abs_ was found to be ∼1.27 at the central wavelength (825 nm) of the OCT source, as shown in Figure 3, which is consistent with the results obtained from OCT phantom imaging. The ratio C_sca_ / C_abs_ suggests that the optical properties of the as-synthesized gold nanocages, for the first time, are dominated by scattering. Compared to previously reported gold nanocages,^11, 15^ the one reported here represents an ∼9-fold increase in the scattering-to-absorption ratio.

**Figure 3.**
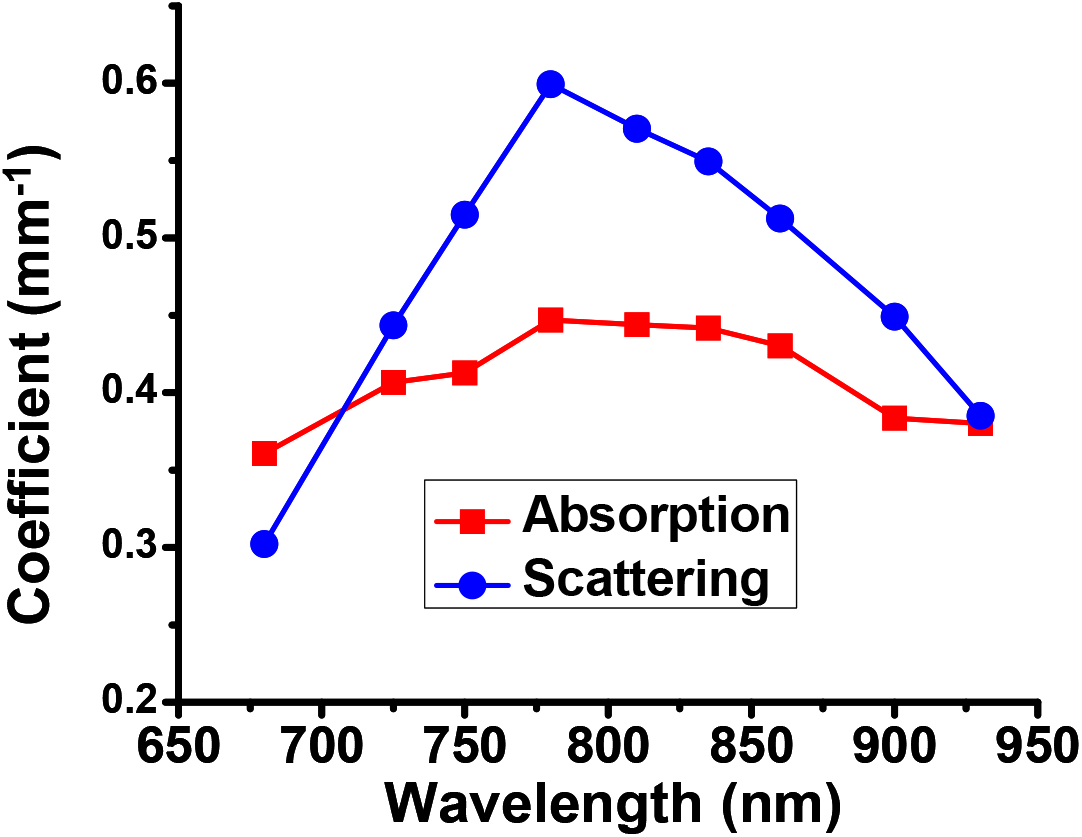
The optical properties of gold nanocages measured by integrating sphere experiments. The results reveal the new gold nanocages are scattering-dominant, with a C_sca_/C_abs_ ratio ∼1.27 at the central wavelength 825 nm of the OCT source.

In addition, we also compared the ratio C_sca_ / C_abs_ of the nanocages with another popular gold nanostructure, gold nanorods. Figure S-1A in the Supporting Information section shows a TEM image of the gold nanorods, which had an aspect ratio of 3.5 and a long axis of ∼46.6 nm. As shown in Figure S-1B, the C_sca_ to C_abs_ ratio of the gold nanorods was ∼0.36, measured by the integrating sphere method at the central wavelength (825 nm) of the OCT source. We find that the newly developed gold nanocages offer an ∼3.5-fold increase in the scattering-to-absorption ratio when compared with nanorods.

To demonstrate the effects of the scattering-dominant gold nanocages on OCT image contrast, we performed OCT imaging of mouse liver and spleen *ex vivo* after tail vein injection of 150 μL of 2 nM PEGylated gold nanocages. The liver and spleen were chosen because these organs quickly capture foreign particles (i.e., the nanocages in this case) after systemic administration. As a control, OCT imaging was also performed on liver and spleen tissues *ex vivo* after a tail vein injection of 150 μL saline. The two pieces of resected liver (or spleen) tissues were placed adjacent to each other, and an OCT image was acquired across both pieces of tissue. Figure 4A shows a combined OCT image of the liver tissues after the injection of saline (on the left, serving as a control) and the injection of PEGylated gold nanocages (on the right), while Figure 4B shows a combined OCT image of the spleen tissues after injection of saline (control) and the injection of PEGylated gold nanocages. We found that the backscattering signals were stronger, with many dark clusters, from the liver and spleen tissues after the administration of gold nanocages compared to the control tissues. These *ex vivo* results demonstrated that the as-synthesized gold nanocages could serve as a scattering contrast agent for OCT imaging, which encouraged us to further investigate the feasibility of the scattering-dominant gold nanocages as a contrast agent for *in vivo* OCT imaging.

**Figure 4.**
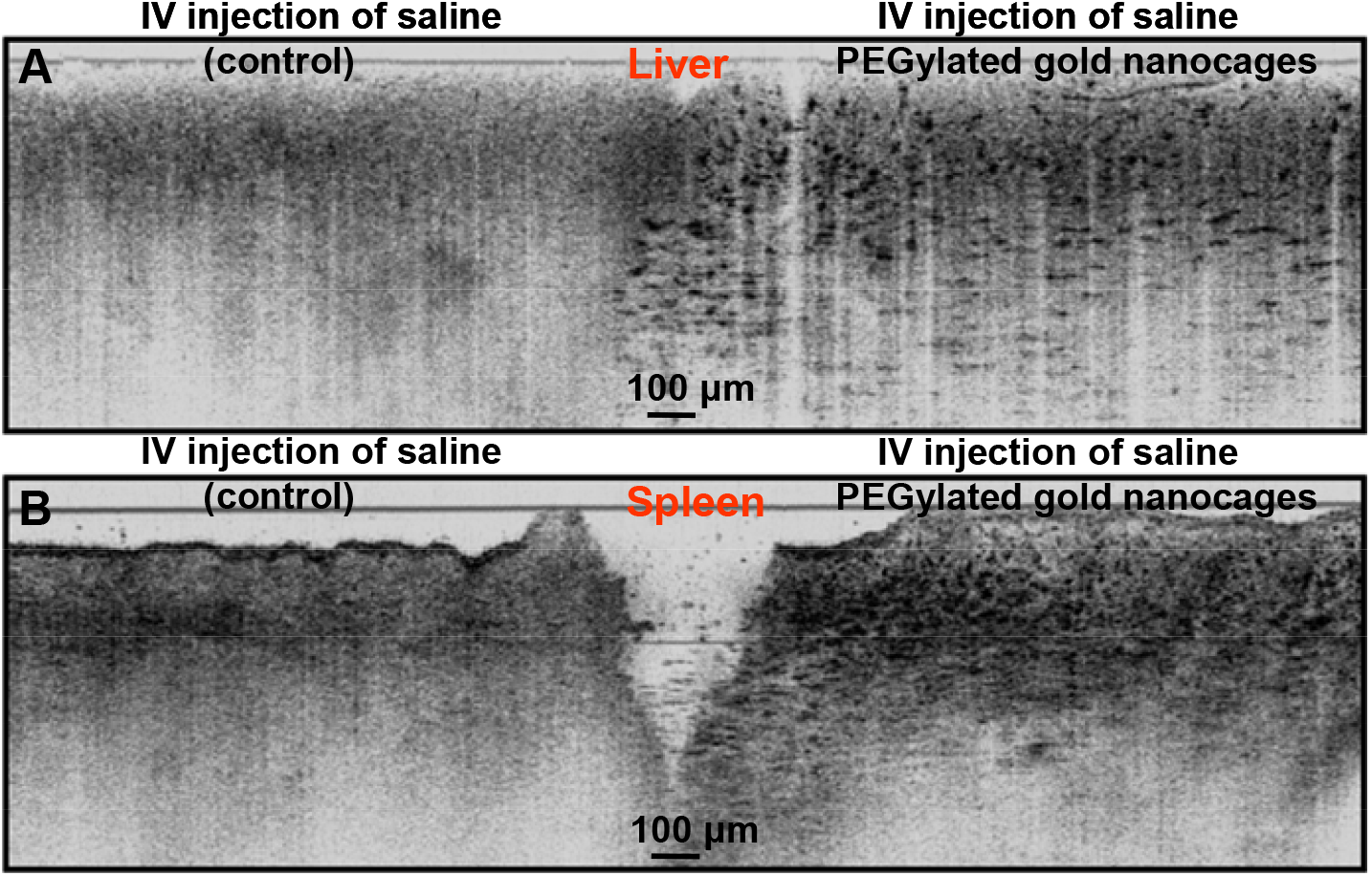
(A) *Ex vivo* OCT image of mouse liver after intravenous injection of saline as control (left) and injection of PEGylated gold nanocages (right). (B) *Ex vivo* OCT image of mouse spleen after intravenous injection of saline as control (left) and injection of PEGylated gold nanocages (right). Stronger backscattered signals were observed from the liver and spleen tissues after intravenous injection of PEGylated gold nanocages.

For *in vivo* experiments, 4 successive tail vein injections of PEGylated gold nanocages (150 μL of 1 nM nanocage aqueous solution per injection) were performed 24 hours apart on a mouse ear tumor model (implanted with the tumor cell line A431). Surface PEGylation, along with the fractionated injection scheme, increased the circulation time of the gold nanocages within the circulatory system and thus enhanced their accumulation in the tumor through the enhanced permeability and retention (EPR) effect.^22, 23^*In vivo* OCT imaging of the mouse tumor was performed 24 hours after the last injection of nanocages. Figures 5A and 5B show a representative OCT image of the tumor before and after the administration of gold nanocages, respectively, and the corresponding OCT intensity decay curves on a logarithmic scale versus imaging depth are shown in Figure 5C. It is evident that the presence of gold nanocages increases the backscattering in the tumor, thus enhancing OCT imaging contrast. The contrast enhancement is approximately 2.4 dB on average, particularly for imaging depths deeper than 400 μm.

**Figure 5.**
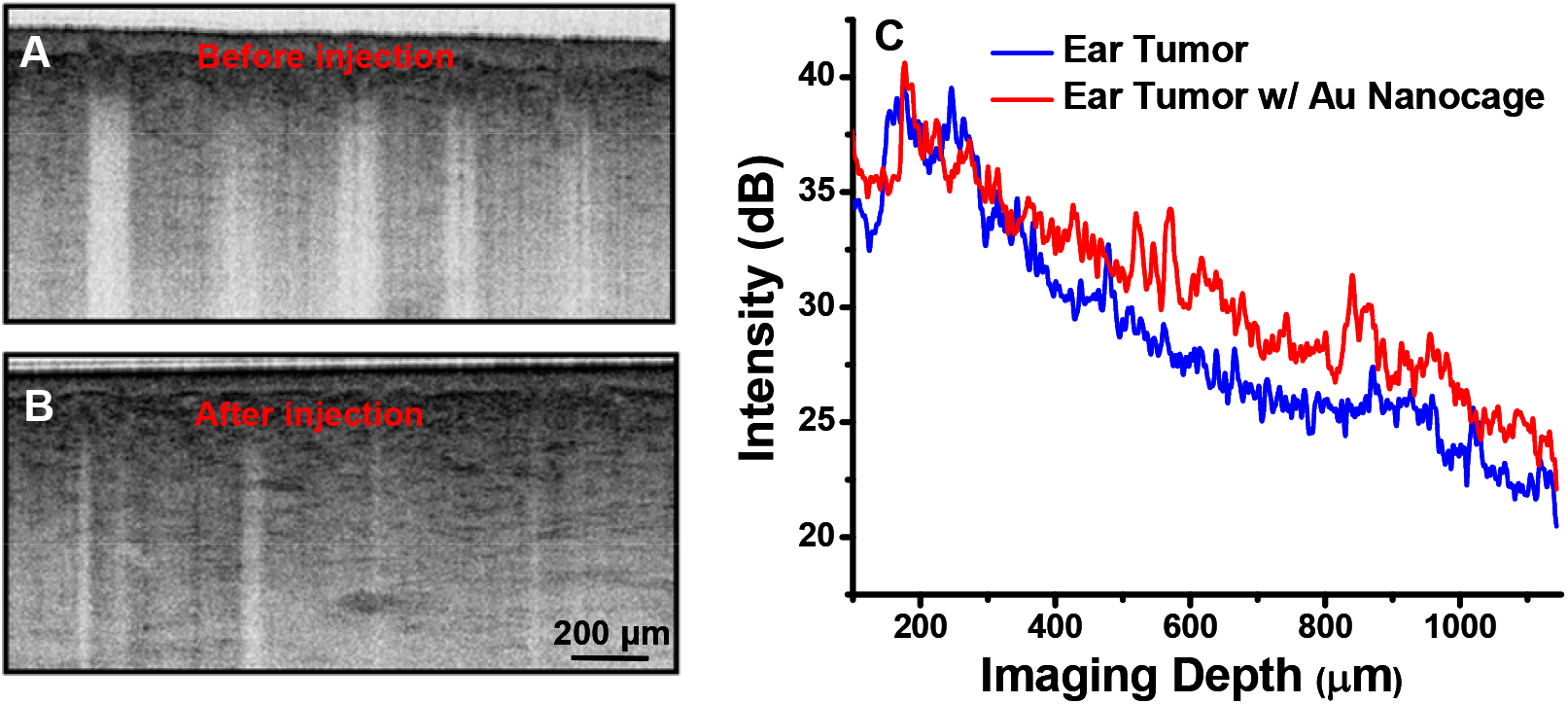
OCT images of a mouse ear tumor (induced with A431 cell line) *in vivo* before (A) and after (B) intravenous injection of gold nanocages. (C) Depth-dependent backscattered OCT intensity plots from a mouse ear tumor on a logarithm scale before (blue curve) and after (red curve) the administration of gold nanocages.

## CONCLUSIONS

In summary, we reported the first scattering-dominant contrast agent based on structured gold nanoparticles (i.e., Au nanocages) to enhance OCT imaging contrast. Unlike previously reported Au nanocages, the new ones were synthesized using a newly developed protocol with a slightly increased size and decreased porosity. This modification led to an approximately 9-fold increase in the ratio of scattering to absorption cross-section. The optical properties of the nanocages were characterized through OCT imaging of nanocage phantoms, with results further validated by integrating sphere measurements. *In vivo* imaging of a tumor model, following systemic administration of the Au nanocages, demonstrated significant contrast enhancement (∼2.4 dB), marking the first successful use of these nanocages for enhanced OCT imaging.

## METHODS

### Synthesis of scattering-dominant gold nanocages with an SPR peak wavelength around 780 nm

Gold nanocages were synthesized using the galvanic replacement reaction between HAuCl_4_ and silver nanocubes.^17^ The basic procedure was similar to the one reported in our previous publications but with some modifications. Briefly, 100 μL of 6 nM aqueous solution of silver nanocube (of ∼ 68 nm in edge length serving as sacrificial templates) was added to 5 mL of deionized water containing PVP (1 mg/mL) in a 50 mL flask under magnetic stirring at room temperature for about 10 minutes. Then, the HAuCl_4_ solution (0.1 mM) was loaded into the flask via syringe pump at a rate of 0.25 mL per minute under magnetic stirring. As the HAuCl_4_ solution was titrated to the flask, the SPR peak of the gold nanocages was monitored by a UV-Vis-NIR spectrophotometer. The titration of the HAuCl_4_ solution was stopped when an appropriate SPR peak was reached and the reaction solution was transferred into a 50 mL centrifuge tube and then centrifuged at 6000 rpm for about 10 minutes. The supernatant was discarded, and the gold nanocages were washed with saturated NaCl solution several times to remove AgCl which was generated during the galvanic replacement reaction. Finally, gold nanocages were washed using deionized water thoroughly and redispersed in 18.1 MΩ-cm E-pure water for further use.

### Quantification of gold nanocage optical properties using OCT phantom images

The depth-dependent OCT backscattered intensity can be described by:

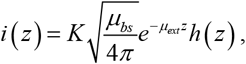

where z is the depth, *μ*_*bs*_ is the backscattering coefficient, *μ*_*ext*_ is the total extinction coefficient, *K* is a system constant and *h*(*z*) is the point-spread function describing the focusing effect of the beam. Since the experimental conditions for both phantoms (with and without gold nanocages) remained the same (i.e., with the same incident power, focused spot size, focusing depth, etc.), the corresponding OCT signals are:

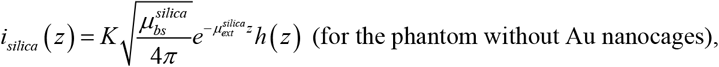

and

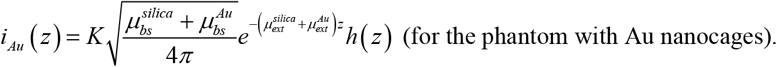

By subtracting the logarithm of the above two signals, we can get a depth (*z*)-dependent curve:

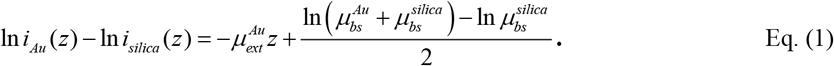

By linearly fitting Eq. (1) versus imaging depth *z*, we are able to get the slope that is the total extinction coefficient 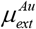 of the gold nanocages. In addition, we will also get the y-axis intercept of the above linear curve (at *z*=0) from the linear fitting, i.e.

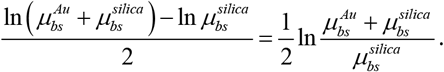

Since the sizes of the silica nanospheres and the gold nanocages are both much smaller than the incident wavelength, the law of Rayleigh scattering can be applied to both nanoparticles. Therefore, the backscattering coefficient should be proportional to the total scattering coefficient for both nanoparticles. Thus we will have:

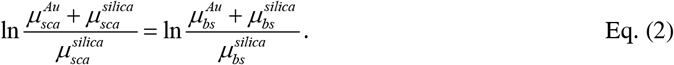

Where 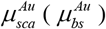 and 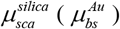 are the total scattering (backscattering) coefficients of the gold nanocages and silica nanospheres, respectively. Furthermore, the total scattering and backscattering coefficients 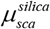 and 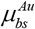 of the silica nanospheres can both be analytically calculated using either Mie or Rayleigh scattering theory. Thus, the total scattering coefficient of the gold nanocages 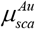 can be derived from Eq. (2). The absorption coefficient of the gold nanocages is then given by 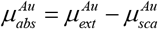.

### Measurement of optical properties of gold nanocages using the integrating sphere method

The optical properties of the gold nanocages were quantitatively characterized by the well-established integrating sphere method in conjunction with a spectrophotometer (Ocean Optics, Dundee, FL). The nanocage solutions (∼0.6 nM) in glass vials were ultrasonicated in a water bath for 20 minutes before the measurements. Custom-made glass cuvettes of 1 mm thickness and 25.5 × 25.5 mm in size were used to hold the solutions during the integrating sphere measurements.

### Gold nanocages as contrast agents for phantom and tissue imaging

OCT imaging was performed on tissue phantoms and mouse tissues with and without the administration of gold nanocages. For phantom experiments, each phantom was made of 5% gelatin embedded with 50 mg/mL silica nanospheres. Gold nanocages were added to one phantom with a final concentration of ∼1 nM. OCT imaging was conducted using a 7-fs Ti:sapphire laser as a light source with a center wavelength at 825 nm and a 3dB spectral bandwidth of 155 nm. As the laser scanned over the two side-by-side phantoms, the OCT intensity signal was measured as a function of imaging depth. For *ex vivo* mouse liver and spleen studies, the mice were sacrificed 24 hours after tail vein administration of 150μL of 2nM PEGylated gold nanocages, followed by immediate tissue resection and OCT imagingAll animal experiments comply with the ARRIVE guidelines and were carried out in accordance with the National Institutes of Health Guide for the Care and Use of Laboratory Animals. All data reported in this study was collected at the Johns Hopkins University (JHU) with approval from JHU Animal Care and Use Committee (under the protocol of MO24M361).

### Gold nanocages as contrast agents for *in vivo* tumor imaging

We examined the feasibility of using the scattering-dominant gold nanocages as a contrast agent for *in vivo* OCT imaging and a mouse xenograft tumor model was used. Male Balb/c nude mice, 6–8 weeks of age and 24.5 g of average weight, were obtained from the Taconic Farmer (One Hudson City Centre, Hudson). Approximately 5 × 10^6^ human epidermoid carcinoma cells (A-431) suspended in 50 μL PBS were injected subcutaneously into the ear of the mice. 10 days after tumor cell inoculation, OCT imaging of the mouse tumor on the ear was performed after 4 fractionated tail vein injections (24 hours apart) of PEGylated Au nanocages (150μL of 1nM solution per injection). Animal experimental procedures in this study were approved by the Institutional Animal Care and Use Committees at the Johns Hopkins University.

## ACKNOWLEDGMENT

This work was supported in part by the National Institutes of Health (NIH) under a grant number R01 CA120480.

## SUPPORTING INFORMATION

**Figure 1-S.**
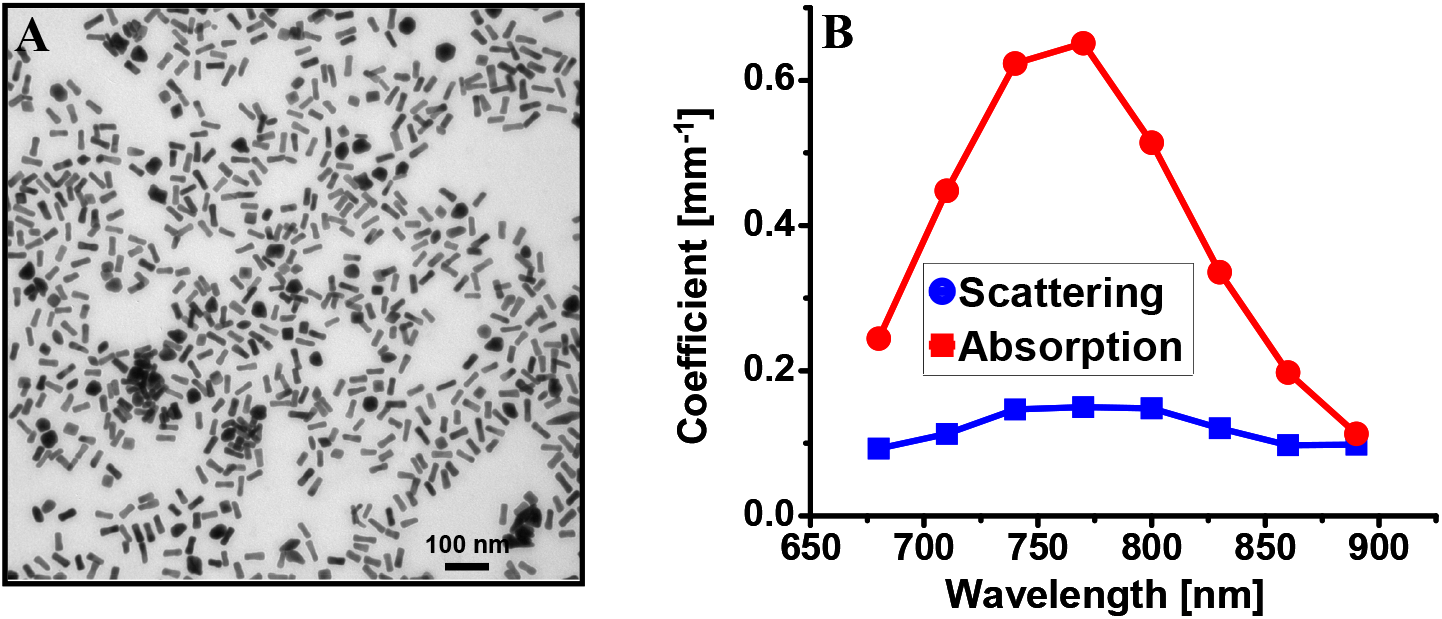
(A) TEM image of gold nanorods of an aspect ratio of 3.5 with a long axis ∼46.6 nm. (B) Integrating sphere measurements reveal gold nanorods are absorption-dominant with a C_sca_/C_abs_ ratio ∼0.36 around the central wavelength of the OCT source (i.e. 825 nm).

